# The Sibship Index can’t discriminates Half-Sibs from Full-Sibs; a new algorithm, including maternal genotype, can do it

**DOI:** 10.1101/265561

**Authors:** Giuseppe Cardillo

**Affiliations:** Viale Colli Aminei, 461 – lotto 29, 80131, Napoli, Italy

## Abstract

Sibship DNA testing is conducted in order to determine if two subjects share one or both biological parents: Full-sibs have both parents in common, whereas Half-sibs have one parent in common, either the mother or the father. When an alleged father is not available for a paternity test, sibship testing is one way to determine family relationships. Results of a sibship test have also been accepted as proof in Social Security benefit and other inheritance claims. To analyze the possibility that the siblings share two common parents versus only one common parent, the ratio of the full sibship index versus half sibship index is computed: a sibship index less than 1, provides genetic evidence which does not favor a full-sibling relationship. In this paper we will demonstrate that the actually used Sibship Index is unable to distinguish Full-sibs from Half-Sibs and the only way to distinguish them is to use a new Index including the genotype of the Mother, that, usually, is the known common parent.

**Highlights:**

- Usually used Sibship Index (SI) greatly overestimates likelihood ratios;
- Rising up the number of investigated STR, all Half-Sibs are declared Full-Sibs;
- The only way to avoid this overestimate is to use a Sibship Index Corrected by the common parent genotype, that usually is the mother (SICMG); all likelihood ratio formulas can be reconducted to five scenarios;
- For both SI and SICMG there is a better cut-off than 1 to discriminate Half-Sibs: this cut-off is computed for all commercial available forensic STR kit.

## Methods

### The likelihood ratio

The probability of sibship after the DNA fingerprint determination is computed using the Bayes’ theorem. To apply the theorem, it is needed to compute, for each analyzed locus, the likelihood ratio: this is the ratio of two maximum probabilities that can be assumed seeing the evidences, under two alternatives hypothesis. In our case, the likelihood ratio is the probability to observe the subjects’ genotypes under the hypothesis they are Full-Sibs over the probability to observe the same genotypes under the hypothesis they are Half-Sibs. Since the loci are independent from each other, the single likelihood ratios can be multiplied together to obtain the cumulative likelihood ratio. The loci used in DNA fingerprint are in Hardy-Weinberg Equilibrium (HWE) and this law will be used to compute the likelihood ratios. Under HWE hypothesis:

- the probability to observe a homozygous locus for the allele A, that has frequency *p* in the population, will be *p*^*2*^;
- the probability to observe a heterozygous locus for the alleles A and B, that have, respectively, frequency *p* and *q* in the population, will be *2pq*;
- the sum of all possible genotypes probabilities is 1.

### Derivation of Sibship Index (SI) for Full-sibs versus Half-sibs

The Sibship Index under the two alternatives hypothesis of Full-sibs and Half-sibs was derived as described elsewhere [1]. Briefly, it can be resumed as follows. For each locus, let *δ*_*ij*_ and *δ*_*kl*_ be the Kronecker’s deltas of the sibs’ alleles:

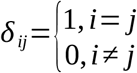

The Kronecker’s delta will be 1 if the locus is homozygous and 0 if the locus is heterozygous. Sibs can share 0, 1 or 2 alleles and so, all possible SI can be resumed using the set theory notation:

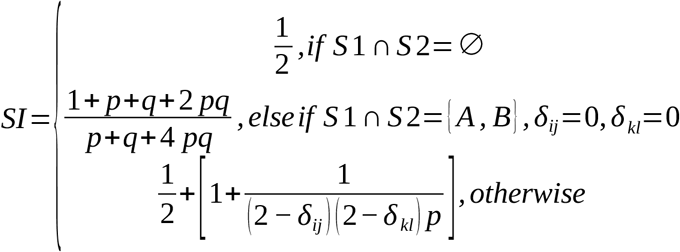

where *p* and *q* are the alleles frequencies in the population. Table 1 summarizes all possible Sibship Index.

**Table 1:** All possible combinations among Mother and Probands and corresponding Sibship Index and Sibship Index Corrected by Maternal Genotype. In Sibship Index, p and q are the frequencies of shared alleles; in the Sibship Index Corrected by Maternal Genotype, p is the frequency of the Paternal shared allele; if Mother and almost one the Sibs are AB then both alleles frequencies are considered.

| Mother |  | Sib 1 |  | Sib 2 |  | Sibship Index | Sibship Index Corrected by Maternal Genotype |
| --- | --- | --- | --- | --- | --- | --- | --- |
| A | A | A | A | A | A | $1/2*(1+1/p)$ | $2/(p*(p+1))$ |
| | | | | A | B | $1/2*(1+1/(2*p))$ | 1/4 |
| | | A | B | A | A | $1/2*(1+1/(2*p))$ | 1/4 |
| | | | | A | B | $(1+p+q+2*p*q)/(p+q+4*p*q)$ | $2/(p*(p+1))$ |
| | | | | A | C | $1/2*(1+1/(4*p))$ | 1/4 |
| A | B | A | A | A | A | $1/2*(1+1/p)$ | $2/(p*(p+1))$ |
| | | | | A | B | $1/2*(1+1/(2*p))$ | $(2*(p+q+1))/(8*q+p*(p+1)^2)$ |
| | | | | A | C | $1/2*(1+1/(2*p))$ | 1/4 |
|  |  |  |  | BB or BC |  | 1/2 | 1/4 |
| | | A | B | AA or BB | | $1/2*(1+1/(2*p))$ | $(2*(p+q+1))/(8*q+p*(p+1)^2)$ |
| | | | | A | B | $(1+p+q+2*p*q)/(p+q+4*p*q)$ | $(2*(p+q)*(p+q+1))/((p*(p+1))^2+(q*(q+1))^2+16*p*q)$ |
| | | | | AC or BC | | $1/2*(1+1/(4*p))$ | 1/4 |
| | | A | C | A | A | $1/2*(1+1/(2*p))$ | 1/4 |
| | | | | AB or AD | | $1/2*(1+1/(4*p))$ | 1/4 |
| | | | | A | C | $(1+p+q+2*p*q)/(p+q+4*p*q)$ | $2/(p*(p+1))$ |
|  |  |  |  | BB or BD |  | 1/2 | 1/4 |
| | | | | B | C | $1/2*(1+1/(4*p))$ | $2/(p*(p+1))$ |

### Derivation of Sibship Index Corrected by Maternal Genotype (SICMG)

There are five possible states for SICMG. In the first one, there are two conditions to verify:

1. maternal and paternal origin of both Sibs alleles can be clearly identifiable because Mother shares only one allele with each of them;
2. Sibs share the paternal allele.

An example of this state is Mother=AB; SIB1=AC and SIB2=BC. To compute the numerator of the SICMG, we assume that Sibs are Full-Sibs. The common Father could be homozygous (*p*^*2*^) and there is 100% of probability he gave the same allele to both Sibs; on the other hand, the Father could be heterozygous (*2pq*) and there is 50% of probability that he gave the allele A to one sib and 50% that he gave the same allele to the other. Using compound and total probability laws:

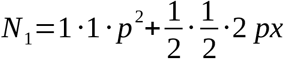

Under the Hardy-Weinberg Equilibrium hypothesis, the sum of all alleles frequencies is equal to 1, so:

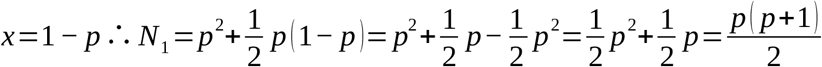

To compute the denominator, we have to hypothesize two different Fathers and each of them could be homozygous or heterozygous, generating four possible combinations of these situations:

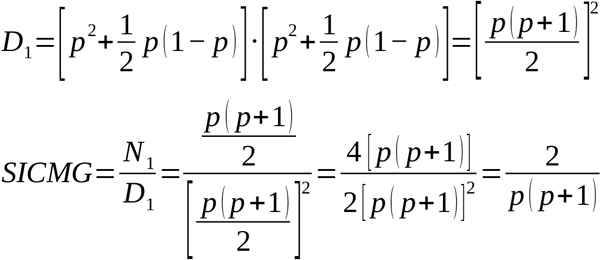

In the second possible state:

1. Again none of the Sibs shares the same heterozygous Mother’s genotype;
2. Sibs do not share the paternal allele.

An example of this state is Mother=AB; SIB1=AC and SIB2=BD. Under the hypothesis that Sibs have the same Father, he can only be heterozygous:

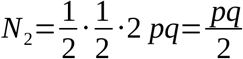

Under the hypothesis of two different Fathers, if one is homozygous or heterozygous for one allele, the other is homozygous or heterozygous for the other allele:

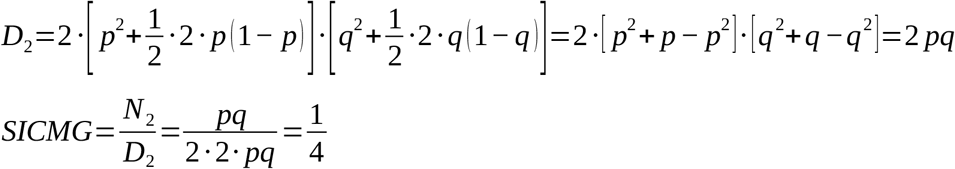

In the third possible state:

1. The Mother is heterozygous and one of the Sibs is heterozygous with the same genotype of the Mother;
2. The other Sib is homozygous.

An example of this state is Mother=AB; SIB1=AB and SIB2=AA. In this case, for the first Sib it is impossible to establish which allele was inherited from the Mother. If the Sibs inherited different maternal alleles, they share the same paternal allele; on the contrary, if they share the same maternal allele, they inherited different paternal alleles.

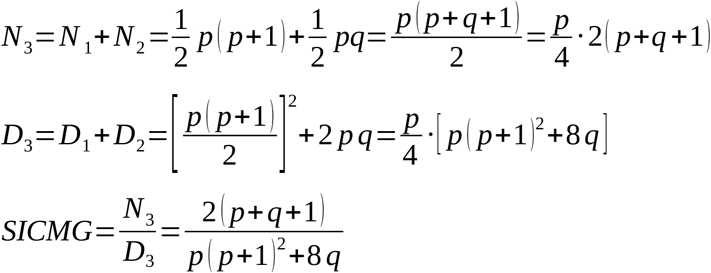

In the fourth possible state:

1. The Mother is heterozygous, one of the Sibs is heterozygous with the same genotype of the Mother;
2. The other Sib is heterozygous and shares only one allele with the first Sib.

An example of this state is Mother=AB; SIB1=AB and SIB2=BC.

This state is equal to the second one, even if the derivation is slightly different.

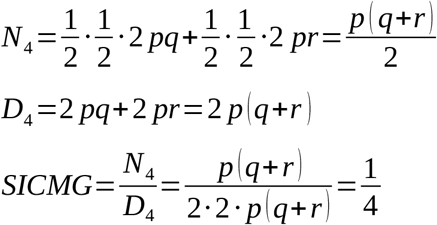

Finally, in the fifth possible state, Mother and Sibs share the same heterozygous genotype: Mother=AB; SIB1=AB and SIB2=AB.

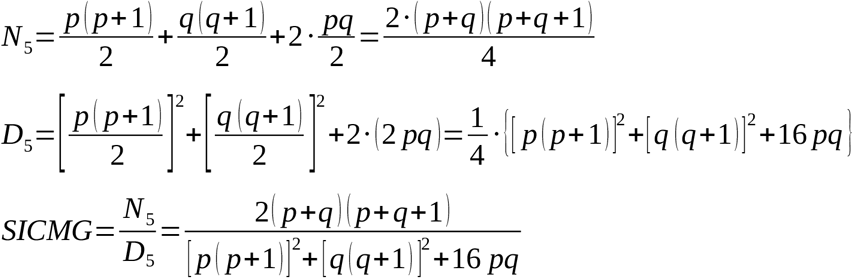

Table 1 summarizes all possible SICMG.

### Algorithm for the simulation of population

To simulate a population, we wrote an algorithm using MatLab R2017a (MathWorks Inc. Natick, MA, USA). We used 50 loci: one half of them are routinely used in commercial kits (CSF1PO, D1S1656, D2S441, D2S1338, D3S1358, D5S818, D6S1043, D7S820, D8S1179, D10S1248, D12S391, D13S317, D16S539, D18S51, D19S433, D21S11, D22S1045, FGA, PENTA D, PENTA E, SE33, TH01, TPOX and vWA); the other is not (D1GATA113, D1S1677, D2S1776, D3S3053, D3S4529, D4S2364, D4S2408, D5S2500, D6S474, D6S1017, D6S1043, D9S1122, D10S1435, D11S4463, D12ATA63, D14S1434, D17S974, D17S1301, D18S853, D20S482, F13A1, F13B, FES, LPL, PENTA B, PENTA C). The Italian allelic frequencies of these loci were retrieved from http://www.allstr.de/allstr/home.seam; for D6S1043, Penta B and Penta C we used the Caucasian frequencies reported by Phillips et al [2]. The input of the algorithm is a column vector of described alleles frequencies for a given locus. In a population it can happen that, for a given locus, some previously described alleles are absent: when this occurs, alleles with null frequency are erased. Matrix multiplication of the column vector by itself transposed generates a square matrix where on the main diagonal there are the squared allelic frequencies (*p*^*2*^ homozygotes) and in the other cells there are products of two allelic frequencies (the matrix is symmetric with respect to the main diagonal). For example:

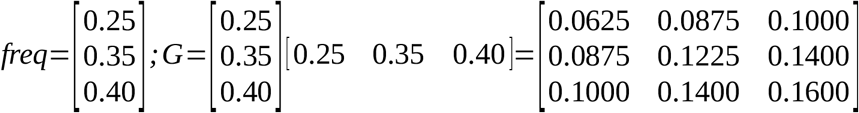

Performing a scalar multiplication for another square matrix that have the same size of the first one, all ones on the main diagonal, all zeros above the main diagonal and all two below the main diagonal, we obtain a matrix where on the main diagonal there are again the *p*^*2*^ homozygotes, below the main diagonal there are the *2pq* heterozygotes and above the main diagonal all values are zero.

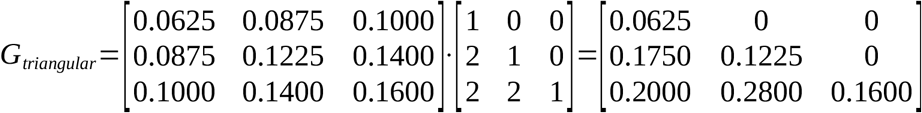

The sum of all matrix elements is 1, as consequence of the Hardy-Weinberg law.

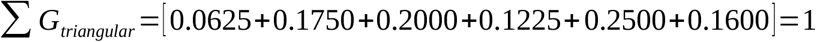

The matrix shows the fraction of the population that has a specific genotype: for example, 6.25% of the population is homozygous for the allele 1 and 17.50% is heterozygous for the alleles 1 and 2. Performing a scalar multiplication for 10 million we obtain a good representation of rare genotypes too:

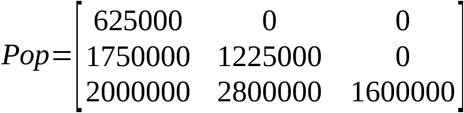

Now, it is possible to construct the genotypes matrix with 10 million rows and 2 columns that represents the genotypes of the population: using this example there are 625000 rows [1 1]; 1750000 rows [1 2] and so on.

The genotypes matrix is randomly shuffled using the Fisher-Yates-Sattolo algorithm and divided in two halves: the first one will be the Mother’s genotypes and the other the Father’s genotypes. We can use these two matrix to generate Full-Sibs pairs.

The mechanism by which the Mother and the Father transmit one or the other of their alleles is like tossing a coin. This process can be simulated using a 5 million rows and 2 columns random matrix chosen from a binomial distribution with parameters p=0.5 and n=1: the first column indicates if from the Mother is inherited the first or the second allele; the second column indicates if from the Father is inherited the first or the second allele. The binomial matrix is reinitialized to generate the second Sib.

To generate Half-Sibs, the Father’s matrix is shuffled again using the Fisher-Yates-Sattolo algorithm and third binomial 5 million rows and 2 columns random matrix is generated. Resuming, there are 5 million pairs of Full-Sibs and 10 million pairs of Half-Sibs. Finally, the algorithm computes the SI and SICMG for each pair of Full-Sibs and Half-Sibs and then stores the results on to the hard disk.

### Determination of single locus and cumulative accuracy of SI and SICMG

SI and SICMG are formulated so that a value greater than 1 suggests that we are observing a Full-Sibs pair and a value lower than 1 suggests that we are observing a Half-Sibs pair. For each locus, we computed how many Full-Sibs had a SI or SICMG greater than 1 and how many Half-Sibs had a SI or SICMG lower than 1. For each locus were determined: Sensitivity, Specificity and Youden’s J Index (JI).The cumulative Index is computed by multiplying together the Indices of single loci. The distributions of SI and SICMG at each locus are not Gaussian but the logarithms of their cumulative distributions are Gaussian, so it is possible to easily compute the crossing points between Full-Sibs and Half-Sibs cumulative distributions obtained from SI and SICMG. In fact, each Gaussian distribution has an equation:

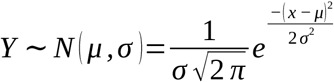

where μ is the mean and σ is the standard deviation. Equating two distribution equations and placing *τ*=1/σ^2^, the crossing points will be found by solving the second degree equation:

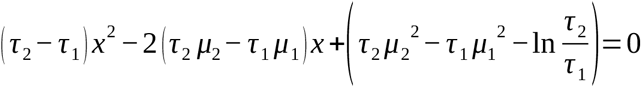

Of course, there are two roots that solve this equation; for our purpose, we used the crossing point (CP) between means. This CP is used to find the Equivalent Error Rate Point (ERRP), that is the point where misclassification rates for both Half-Sibs and Full-Sibs are equal (Sensitivity=Specificity). To find ERRP we used Bisection Method using CP as guessing starting point to find the zero of the function:

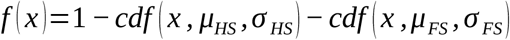

where the Cumulative Density Function (cdf) for a Gaussian distribution is:

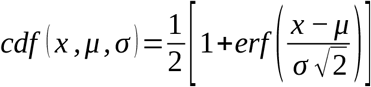

and the Error Function (erf) is:

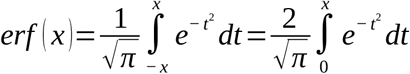

Again, the performances of cumulative SI and SICMG were computed using as threshold 1 or the ERRP.

## Results and Discussion

Tables 2 and 3 summarize, respectively, SI and SICMG performances computed for each locus and cumulatively. STRs are sorted to maximize the Youden’s J index (JI). A randomization t-test [3] using a Monte Carlo naïve method, performed on the arrays of JI showed a statistically significative difference between methods (p-value=0). Moreover, when SI algorithm is used, increasing the number of considered loci, JI dramatically decreases (Table 2); on the contrary, when SICMG algorithm is used JI increases (Table 3). It is interesting to highlight that, when SICMG algorithm is used, the JI increases, reaches a maximum and then decreases (Table 3). In the sibship analysis, usually, a likelihood ratio greater than 1 indicates a Full-Sibs kinship and a likelihood ratio lower than 1 indicates a Half-Sibs kinship. As shown in Table 2, when all loci are used to compute SI, all couples have a likelihood ratio grater than 1 and so the SI is completely useless to discriminate Half-Sibs from Full-Sibs. The phenomenon is clearly highlighted in Figure 1: when unusual STRs are added and SI algorithm is used, both Full-Sibs and Half-Sibs distributions means move to the right and about 35% of the areas overlaps; on the contrary, when SICMG algorithm is used, Full-Sibs distribution mean moves to the right, the Half-Sibs distribution mean moves to the left and the areas overlap is about 3%, ten fold lower than SI overlap.

**Table 2:**
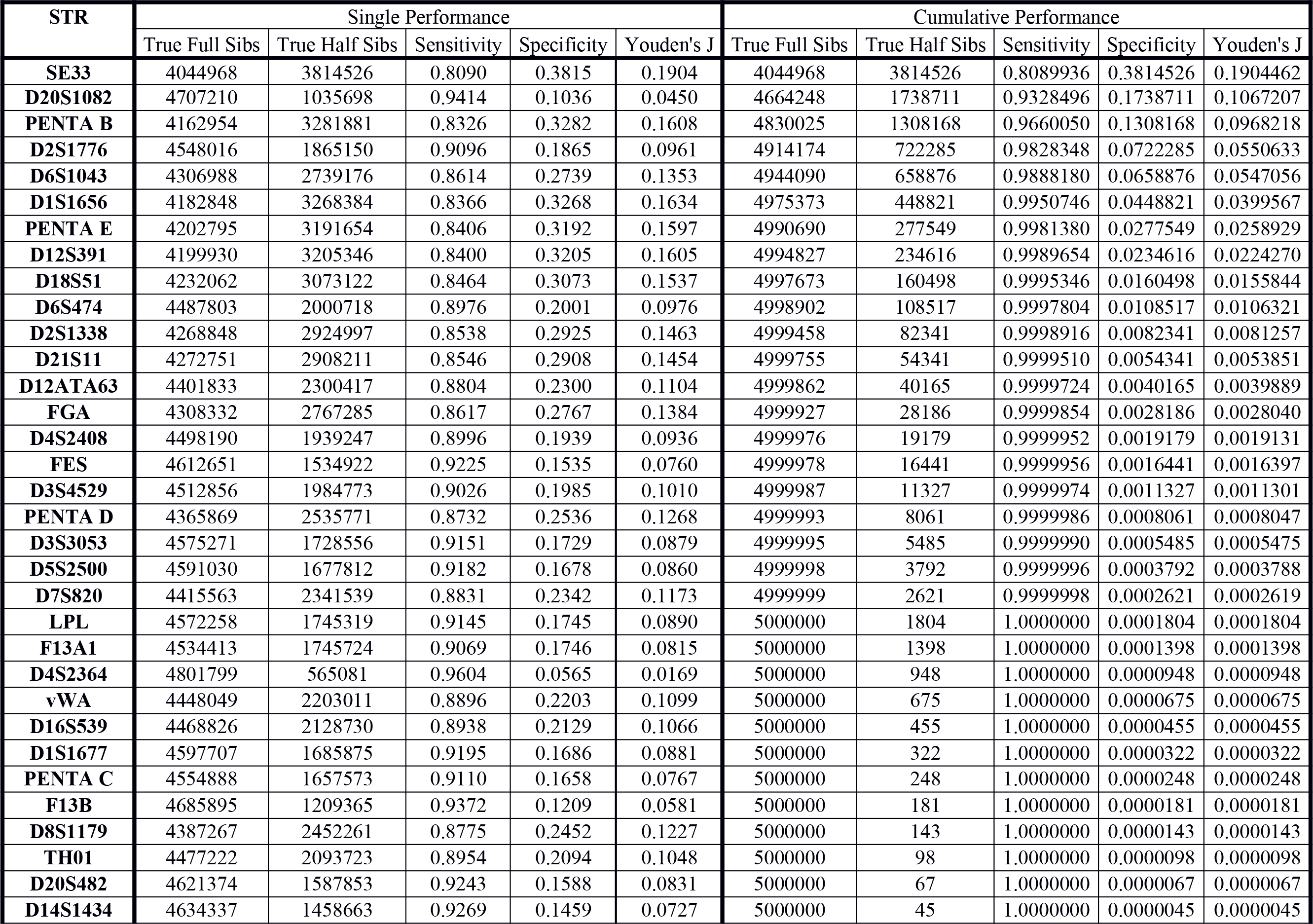
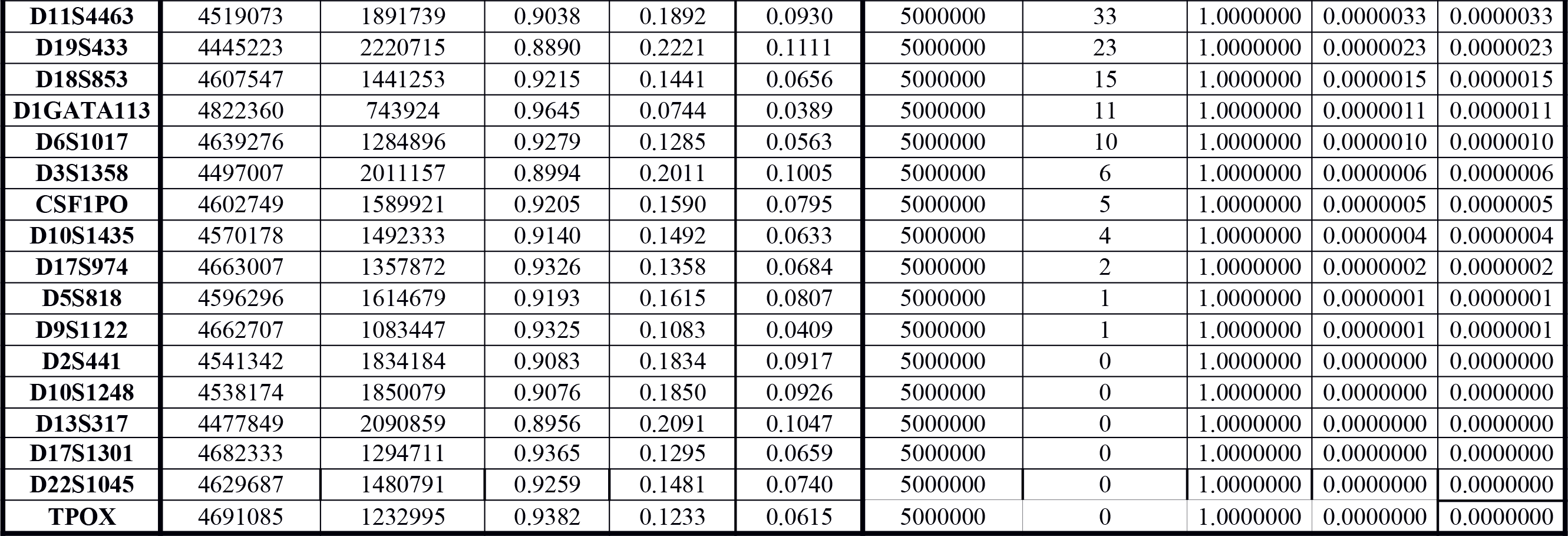
Performance of the Sibship Index (SI) for each locus and cumulatively computed.

| STR | Single Performance |  |  |  |  | Cumulative Performance |  |  |  |  |
| --- | --- | --- | --- | --- | --- | --- | --- | --- | --- | --- |
|  | True Full Sibs | True Half Sibs | Sensitivity | Specificity | Youden's J | True Full Sibs | True Half Sibs | Sensitivity | Specificity | Youden's J |
| SE33 | 4044968 | 3814526 | 0.8090 | 0.3815 | 0.1904 | 4044968 | 3814526 | 0.8089936 | 0.3814526 | 0.1904462 |
| D20S1082 | 4707210 | 1035698 | 0.9414 | 0.1036 | 0.0450 | 4664248 | 1738711 | 0.9328496 | 0.1738711 | 0.1067207 |
| PENTA B | 4162954 | 3281881 | 0.8326 | 0.3282 | 0.1608 | 4830025 | 1308168 | 0.9660050 | 0.1308168 | 0.0968218 |
| D2S1776 | 4548016 | 1865150 | 0.9096 | 0.1865 | 0.0961 | 4914174 | 722285 | 0.9828348 | 0.0722285 | 0.0550633 |
| D6S1043 | 4306988 | 2739176 | 0.8614 | 0.2739 | 0.1353 | 4944090 | 658876 | 0.9888180 | 0.0658876 | 0.0547056 |
| D1S1656 | 4182848 | 3268384 | 0.8366 | 0.3268 | 0.1634 | 4975373 | 448821 | 0.9950746 | 0.0448821 | 0.0399567 |
| PENTA E | 4202795 | 3191654 | 0.8406 | 0.3192 | 0.1597 | 4990690 | 277549 | 0.9981380 | 0.0277549 | 0.0258929 |
| D12S391 | 4199930 | 3205346 | 0.8400 | 0.3205 | 0.1605 | 4994827 | 234616 | 0.9989654 | 0.0234616 | 0.0224270 |
| D18S51 | 4232062 | 3073122 | 0.8464 | 0.3073 | 0.1537 | 4997673 | 160498 | 0.9995346 | 0.0160498 | 0.0155844 |
| D6S474 | 4487803 | 2000718 | 0.8976 | 0.2001 | 0.0976 | 4998902 | 108517 | 0.9997804 | 0.0108517 | 0.0106321 |
| D2S1338 | 4268848 | 2924997 | 0.8538 | 0.2925 | 0.1463 | 4999458 | 82341 | 0.9998916 | 0.0082341 | 0.0081257 |
| D21S11 | 4272751 | 2908211 | 0.8546 | 0.2908 | 0.1454 | 4999755 | 54341 | 0.9999510 | 0.0054341 | 0.0053851 |
| D12ATA63 | 4401833 | 2300417 | 0.8804 | 0.2300 | 0.1104 | 4999862 | 40165 | 0.9999724 | 0.0040165 | 0.0039889 |
| FGA | 4308332 | 2767285 | 0.8617 | 0.2767 | 0.1384 | 4999927 | 28186 | 0.9999854 | 0.0028186 | 0.0028040 |
| D4S2408 | 4498190 | 1939247 | 0.8996 | 0.1939 | 0.0936 | 4999976 | 19179 | 0.9999952 | 0.0019179 | 0.0019131 |
| FES | 4612651 | 1534922 | 0.9225 | 0.1535 | 0.0760 | 4999978 | 16441 | 0.9999956 | 0.0016441 | 0.0016397 |
| D3S4529 | 4512856 | 1984773 | 0.9026 | 0.1985 | 0.1010 | 4999987 | 11327 | 0.9999974 | 0.0011327 | 0.0011301 |
| PENTA D | 4365869 | 2535771 | 0.8732 | 0.2536 | 0.1268 | 4999993 | 8061 | 0.9999986 | 0.0008061 | 0.0008047 |
| D3S3053 | 4575271 | 1728556 | 0.9151 | 0.1729 | 0.0879 | 4999995 | 5485 | 0.9999990 | 0.0005485 | 0.0005475 |
| D5S2500 | 4591030 | 1677812 | 0.9182 | 0.1678 | 0.0860 | 4999998 | 3792 | 0.9999996 | 0.0003792 | 0.0003788 |
| D7S820 | 4415563 | 2341539 | 0.8831 | 0.2342 | 0.1173 | 4999999 | 2621 | 0.9999998 | 0.0002621 | 0.0002619 |
| LPL | 4572258 | 1745319 | 0.9145 | 0.1745 | 0.0890 | 5000000 | 1804 | 1.0000000 | 0.0001804 | 0.0001804 |
| F13A1 | 4534413 | 1745724 | 0.9069 | 0.1746 | 0.0815 | 5000000 | 1398 | 1.0000000 | 0.0001398 | 0.0001398 |
| D4S2364 | 4801799 | 565081 | 0.9604 | 0.0565 | 0.0169 | 5000000 | 948 | 1.0000000 | 0.0000948 | 0.0000948 |
| vWA | 4448049 | 2203011 | 0.8896 | 0.2203 | 0.1099 | 5000000 | 675 | 1.0000000 | 0.0000675 | 0.0000675 |
| D16S539 | 4468826 | 2128730 | 0.8938 | 0.2129 | 0.1066 | 5000000 | 455 | 1.0000000 | 0.0000455 | 0.0000455 |
| D1S1677 | 4597707 | 1685875 | 0.9195 | 0.1686 | 0.0881 | 5000000 | 322 | 1.0000000 | 0.0000322 | 0.0000322 |
| PENTA C | 4554888 | 1657573 | 0.9110 | 0.1658 | 0.0767 | 5000000 | 248 | 1.0000000 | 0.0000248 | 0.0000248 |
| F13B | 4685895 | 1209365 | 0.9372 | 0.1209 | 0.0581 | 5000000 | 181 | 1.0000000 | 0.0000181 | 0.0000181 |
| D8S1179 | 4387267 | 2452261 | 0.8775 | 0.2452 | 0.1227 | 5000000 | 143 | 1.0000000 | 0.0000143 | 0.0000143 |
| TH01 | 4477222 | 2093723 | 0.8954 | 0.2094 | 0.1048 | 5000000 | 98 | 1.0000000 | 0.0000098 | 0.0000098 |
| D20S482 | 4621374 | 1587853 | 0.9243 | 0.1588 | 0.0831 | 5000000 | 67 | 1.0000000 | 0.0000067 | 0.0000067 |
| D14S1434 | 4634337 | 1458663 | 0.9269 | 0.1459 | 0.0727 | 5000000 | 45 | 1.0000000 | 0.0000045 | 0.0000045 |
| **D11S4463** | 4519073 | 1891739 | 0.9038 | 0.1892 | 0.0930 | 5000000 | 33 | 1.0000000 | 0.0000033 | 0.0000033 |
| **D19S433** | 4445223 | 2220715 | 0.8890 | 0.2221 | 0.1111 | 5000000 | 23 | 1.0000000 | 0.0000023 | 0.0000023 |
| **D18S853** | 4607547 | 1441253 | 0.9215 | 0.1441 | 0.0656 | 5000000 | 15 | 1.0000000 | 0.0000015 | 0.0000015 |
| **D1GATA113** | 4822360 | 743924 | 0.9645 | 0.0744 | 0.0389 | 5000000 | 11 | 1.0000000 | 0.0000011 | 0.0000011 |
| **D6S1017** | 4639276 | 1284896 | 0.9279 | 0.1285 | 0.0563 | 5000000 | 10 | 1.0000000 | 0.0000010 | 0.0000010 |
| **D3S1358** | 4497007 | 2011157 | 0.8994 | 0.2011 | 0.1005 | 5000000 | 6 | 1.0000000 | 0.0000006 | 0.0000006 |
| **CSF1PO** | 4602749 | 1589921 | 0.9205 | 0.1590 | 0.0795 | 5000000 | 5 | 1.0000000 | 0.0000005 | 0.0000005 |
| **D10S1435** | 4570178 | 1492333 | 0.9140 | 0.1492 | 0.0633 | 5000000 | 4 | 1.0000000 | 0.0000004 | 0.0000004 |
| **D17S974** | 4663007 | 1357872 | 0.9326 | 0.1358 | 0.0684 | 5000000 | 2 | 1.0000000 | 0.0000002 | 0.0000002 |
| **D5S818** | 4596296 | 1614679 | 0.9193 | 0.1615 | 0.0807 | 5000000 | 1 | 1.0000000 | 0.0000001 | 0.0000001 |
| **D9S1122** | 4662707 | 1083447 | 0.9325 | 0.1083 | 0.0409 | 5000000 | 1 | 1.0000000 | 0.0000001 | 0.0000001 |
| **D2S441** | 4541342 | 1834184 | 0.9083 | 0.1834 | 0.0917 | 5000000 | 0 | 1.0000000 | 0.0000000 | 0.0000000 |
| **D10S1248** | 4538174 | 1850079 | 0.9076 | 0.1850 | 0.0926 | 5000000 | 0 | 1.0000000 | 0.0000000 | 0.0000000 |
| **D13S317** | 4477849 | 2090859 | 0.8956 | 0.2091 | 0.1047 | 5000000 | 0 | 1.0000000 | 0.0000000 | 0.0000000 |
| **D17S1301** | 4682333 | 1294711 | 0.9365 | 0.1295 | 0.0659 | 5000000 | 0 | 1.0000000 | 0.0000000 | 0.0000000 |
| **D22S1045** | 4629687 | 1480791 | 0.9259 | 0.1481 | 0.0740 | 5000000 | 0 | 1.0000000 | 0.0000000 | 0.0000000 |
| **TPOX** | 4691085 | 1232995 | 0.9382 | 0.1233 | 0.0615 | 5000000 | 0 | 1.0000000 | 0.0000000 | 0.0000000 |

**Table 3:** Performance of the Sibship Index Corrected by Maternal Genotype (SICMG) for each locus and cumulatively computed.

| STR | Single Performance |  |  |  |  | Cumulative Performance |  |  |  |  |
| --- | --- | --- | --- | --- | --- | --- | --- | --- | --- | --- |
|  | True Full Sibs | True Half Sibs | Sensitivity | Specificity | Youden's J | True Full Sibs | True Half Sibs | Sensitivity | Specificity | Youden's J |
| SE33 | 2591958 | 9318668 | 0.5184 | 0.9319 | 0.4503 | 2591958 | 9318668 | 0.5183916 | 0.9318668 | 0.4502584 |
| D1S1656 | 2652355 | 8933171 | 0.5305 | 0.8933 | 0.4238 | 3712974 | 8535908 | 0.7425948 | 0.8535908 | 0.5961856 |
| D12S391 | 2663306 | 8882135 | 0.5327 | 0.8882 | 0.4209 | 3730809 | 8770834 | 0.7461618 | 0.8770834 | 0.6232452 |
| PENTA E | 2665605 | 8867483 | 0.5331 | 0.8867 | 0.4199 | 3713086 | 9456691 | 0.7426172 | 0.9456691 | 0.6882863 |
| D18S51 | 2680617 | 8777175 | 0.5361 | 0.8777 | 0.4138 | 4052961 | 9340945 | 0.8105922 | 0.9340945 | 0.7446867 |
| PENTA B | 2739600 | 8468239 | 0.5479 | 0.8468 | 0.3947 | 4071341 | 9514549 | 0.8142682 | 0.9514549 | 0.7657231 |
| D21S11 | 2705720 | 8626140 | 0.5411 | 0.8626 | 0.4038 | 4162829 | 9641678 | 0.8325658 | 0.9641678 | 0.7967336 |
| D2S1338 | 2702576 | 8646281 | 0.5405 | 0.8646 | 0.4051 | 4280818 | 9651741 | 0.8561636 | 0.9651741 | 0.8213377 |
| D6S1043 | 2838203 | 8119458 | 0.5676 | 0.8119 | 0.3796 | 4326337 | 9727393 | 0.8652674 | 0.9727393 | 0.8380067 |
| FGA | 2719592 | 8516592 | 0.5439 | 0.8517 | 0.3956 | 4379815 | 9777702 | 0.8759630 | 0.9777702 | 0.8537332 |
| PENTA D | 2758541 | 8301877 | 0.5517 | 0.8302 | 0.3819 | 4422081 | 9808087 | 0.8844162 | 0.9808087 | 0.8652249 |
| D8S1179 | 2777374 | 8210247 | 0.5555 | 0.8210 | 0.3765 | 4448568 | 9850096 | 0.8897136 | 0.9850096 | 0.8747232 |
| D19S433 | 2830801 | 7938249 | 0.5662 | 0.7938 | 0.3600 | 4481349 | 9871149 | 0.8962698 | 0.9871149 | 0.8833847 |
| D12ATA63 | 2948278 | 7282833 | 0.5897 | 0.7283 | 0.3179 | 4514045 | 9880568 | 0.9028090 | 0.9880568 | 0.8908658 |
| D7S820 | 2795659 | 8104051 | 0.5591 | 0.8104 | 0.3695 | 4537756 | 9899704 | 0.9075512 | 0.9899704 | 0.8975216 |
| D3S4529 | 2941294 | 7487810 | 0.5883 | 0.7488 | 0.3370 | 4563995 | 9906242 | 0.9127990 | 0.9906242 | 0.9034232 |
| D6S474 | 3021899 | 7110574 | 0.6044 | 0.7111 | 0.3154 | 4589089 | 9910202 | 0.9178178 | 0.9910202 | 0.9088380 |
| vWA | 2822565 | 7944315 | 0.5645 | 0.7944 | 0.3589 | 4606657 | 9924019 | 0.9213314 | 0.9924019 | 0.9137333 |
| D4S2408 | 3000494 | 7138532 | 0.6001 | 0.7139 | 0.3140 | 4626495 | 9927698 | 0.9252990 | 0.9927698 | 0.9180688 |
| D16S539 | 2845411 | 7848506 | 0.5691 | 0.7849 | 0.3539 | 4641622 | 9938740 | 0.9283244 | 0.9938740 | 0.9221984 |
| D13S317 | 2852159 | 7802579 | 0.5704 | 0.7803 | 0.3507 | 4655335 | 9948290 | 0.9310670 | 0.9948290 | 0.9258960 |
| D3S3053 | 3159246 | 6559247 | 0.6318 | 0.6559 | 0.2878 | 4670354 | 9950273 | 0.9340708 | 0.9950273 | 0.9290981 |
| D2S441 | 2923072 | 7457641 | 0.5846 | 0.7458 | 0.3304 | 4681215 | 9957676 | 0.9362430 | 0.9957676 | 0.9320106 |
| TH01 | 2843199 | 7837043 | 0.5686 | 0.7837 | 0.3523 | 4691802 | 9964150 | 0.9383604 | 0.9964150 | 0.9347754 |
| D3S1358 | 2862199 | 7735336 | 0.5724 | 0.7735 | 0.3460 | 4701286 | 9968979 | 0.9402572 | 0.9968979 | 0.9371551 |
| D5S2500 | 3134986 | 6700703 | 0.6270 | 0.6701 | 0.2971 | 4712176 | 9970293 | 0.9424352 | 0.9970293 | 0.9394645 |
| D11S4463 | 2985294 | 6928844 | 0.5971 | 0.6929 | 0.2899 | 4722928 | 9971284 | 0.9445856 | 0.9971284 | 0.9417140 |
| D10S1248 | 2910053 | 7519876 | 0.5820 | 0.7520 | 0.3340 | 4730143 | 9975497 | 0.9460286 | 0.9975497 | 0.9435783 |
| LPL | 3044556 | 6908648 | 0.6089 | 0.6909 | 0.2998 | 4738126 | 9976650 | 0.9476252 | 0.9976650 | 0.9452902 |
| D14S1434 | 3279335 | 5934420 | 0.6559 | 0.5934 | 0.2493 | 4745040 | 9976763 | 0.9490080 | 0.9976763 | 0.9466843 |
| **D5S818** | 2775334 | 7466223 | 0.5551 | 0.7466 | 0.3017 | 4750327 | 9979865 | 0.9500654 | 0.9979865 | 0.9480519 |
| **D18S853** | 2652423 | 7522221 | 0.5305 | 0.7522 | 0.2827 | 4756342 | 9980916 | 0.9512684 | 0.9980916 | 0.9493600 |
| **D22S1045** | 2539991 | 7712817 | 0.5080 | 0.7713 | 0.2793 | 4760439 | 9983358 | 0.9520878 | 0.9983358 | 0.9504236 |
| **D1S1677** | 3043564 | 6836021 | 0.6087 | 0.6836 | 0.2923 | 4765268 | 9984465 | 0.9530536 | 0.9984465 | 0.9515001 |
| **CSF1PO** | 2981845 | 7131608 | 0.5964 | 0.7132 | 0.3095 | 4768968 | 9986499 | 0.9537936 | 0.9986499 | 0.9524435 |
| **D20S482** | 3164263 | 6369768 | 0.6329 | 0.6370 | 0.2698 | 4772546 | 9987060 | 0.9545092 | 0.9987060 | 0.9532152 |
| **TPOX** | 2946662 | 6797909 | 0.5893 | 0.6798 | 0.2691 | 4775382 | 9988608 | 0.9550764 | 0.9988608 | 0.9539372 |
| **D10S1435** | 2698804 | 7285156 | 0.5398 | 0.7285 | 0.2683 | 4777904 | 9989130 | 0.9555808 | 0.9989130 | 0.9544938 |
| **D20S1082** | 2663714 | 7280251 | 0.5327 | 0.7280 | 0.2608 | 4779053 | 9989576 | 0.9558106 | 0.9989576 | 0.9547682 |
| **D17S974** | 2545028 | 7388159 | 0.5090 | 0.7388 | 0.2478 | 4779525 | 9990089 | 0.9559050 | 0.9990089 | 0.9549139 |
| **D6S1017** | 2917211 | 6803358 | 0.5834 | 0.6803 | 0.2638 | 4780490 | 9990718 | 0.9560980 | 0.9990718 | 0.9551698 |
| **D17S1301** | 2379392 | 7610555 | 0.4759 | 0.7611 | 0.2369 | 4780656 | 9991175 | 0.9561312 | 0.9991175 | 0.9552487 |
| **PENTA C** | 3045620 | 6624301 | 0.6091 | 0.6624 | 0.2716 | 4781237 | 9991031 | 0.9562474 | 0.9991031 | 0.9553505 |
| **D2S1776** | 3162890 | 6537791 | 0.6326 | 0.6538 | 0.2864 | 4778311 | 9990991 | 0.9556622 | 0.9990991 | 0.9547613 |
| **D9S1122** | 2999529 | 5902130 | 0.5999 | 0.5902 | 0.1901 | 4775210 | 9990974 | 0.9550420 | 0.9990974 | 0.9541394 |
| **D1GATA113** | 3581165 | 4157776 | 0.7162 | 0.4158 | 0.1320 | 4771850 | 9991185 | 0.9543700 | 0.9991185 | 0.9534885 |
| **F13A1** | 3105796 | 6253732 | 0.6212 | 0.6254 | 0.2465 | 4769291 | 9990387 | 0.9538582 | 0.9990387 | 0.9528969 |
| **FES** | 3195207 | 6340255 | 0.6390 | 0.6340 | 0.2731 | 4765215 | 9990034 | 0.9530430 | 0.9990034 | 0.9520464 |
| **F13B** | 3299071 | 5448406 | 0.6598 | 0.5448 | 0.2047 | 4758536 | 9989157 | 0.9517072 | 0.9989157 | 0.9506229 |
| **D4S2364** | 3053343 | 5296001 | 0.6107 | 0.5296 | 0.1403 | 4740958 | 9987558 | 0.9481916 | 0.9987558 | 0.9469474 |

**Figure 1:**
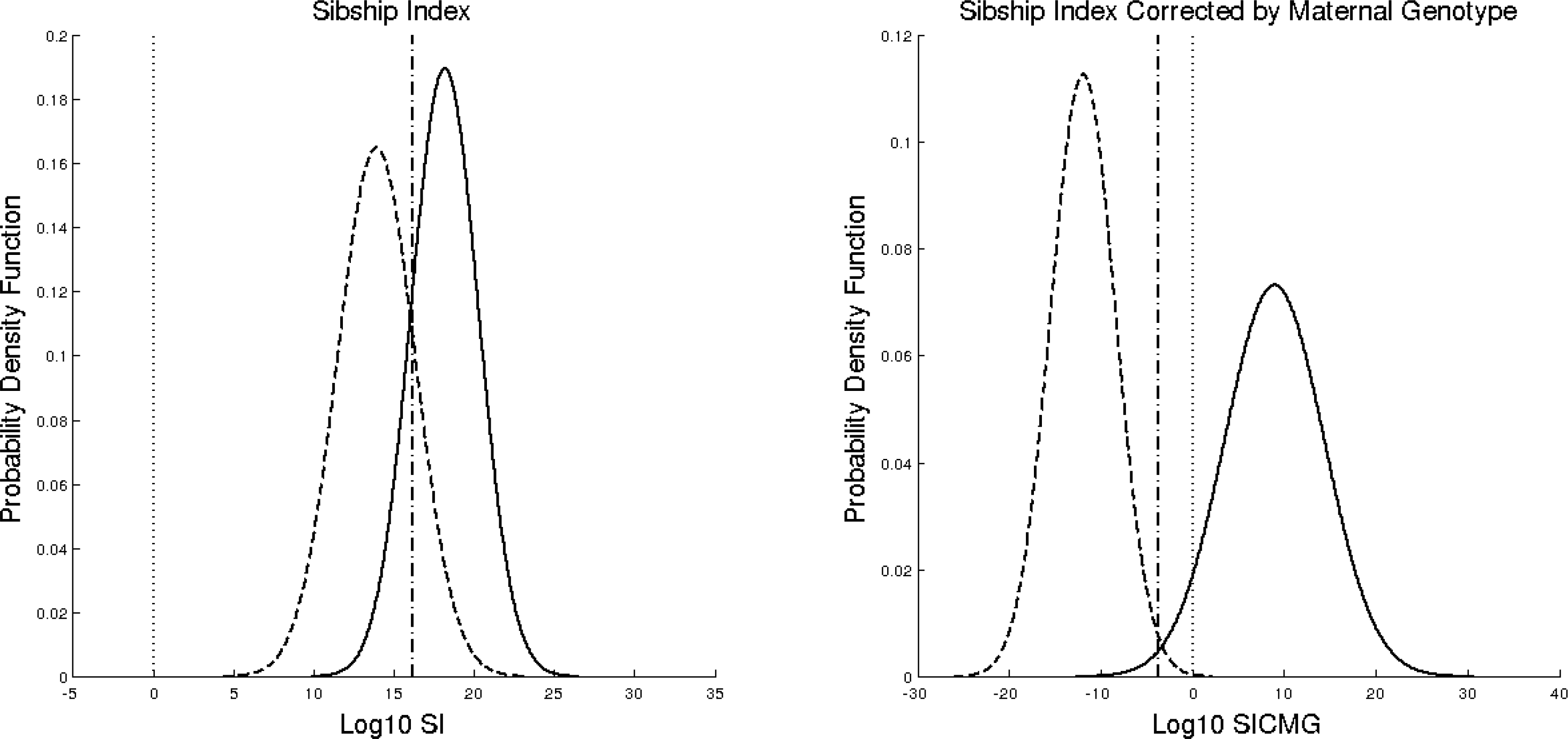
Distributions of logarithms of cumulative Sibship Index and Sibship Index Corrected by Maternal Genotype. Solid line = Full Sibs; dashed line = Half Sibs; dotted line = Cumulative Likelihood Ratio cut off set at 1; dash dot line = Cumulative Likelihood Ratio cut off set as Equivalent Error Rate Point.

To improve the results, the EERP must be taken in account and Table 4 shows the performances of commercial kits.

**Table 4:** Performances of different commercial available test in Full-Sibs versus Half-Sibs discrimination using Sibship Index corrected by maternal genotype. 1 = the STR is present in the kit; 0 = the STR is absent in the kit.

| STR | LIFE Technologies - AmpFLSTR |  |  |  | Promega – Powerplex |  |  |  |  | CORE |
| --- | --- | --- | --- | --- | --- | --- | --- | --- | --- | --- |
|  | Identifiler | NGM | NGM Select | GlobalFiler | 16 | 18D | 17ESX | 21 | Fusion |  |
| CSF1PO | 1 | 0 | 0 | 1 | 1 | 1 | 0 | 1 | 1 | 1 |
| D1S1656 | 0 | 1 | 1 | 1 | 0 | 0 | 1 | 1 | 1 | 1 |
| D2S441 | 0 | 1 | 1 | 1 | 0 | 1 | 1 | 0 | 1 | 1 |
| D2S1338 | 1 | 1 | 1 | 1 | 0 | 0 | 1 | 1 | 1 | 1 |
| D3S1358 | 1 | 1 | 1 | 1 | 1 | 1 | 1 | 1 | 1 | 1 |
| D5S818 | 1 | 0 | 0 | 1 | 1 | 1 | 0 | 1 | 1 | 1 |
| D6S1043 | 0 | 0 | 0 | 0 | 0 | 0 | 0 | 1 | 0 | 1 |
| D7S820 | 1 | 0 | 0 | 1 | 1 | 1 | 0 | 1 | 1 | 1 |
| D8S1179 | 1 | 1 | 1 | 1 | 1 | 1 | 1 | 1 | 1 | 1 |
| D10S1248 | 0 | 1 | 1 | 1 | 0 | 0 | 1 | 0 | 1 | 1 |
| D12S391 | 0 | 1 | 1 | 1 | 0 | 0 | 1 | 1 | 1 | 1 |
| D13S317 | 1 | 0 | 0 | 1 | 1 | 1 | 0 | 1 | 1 | 1 |
| D16S539 | 1 | 1 | 1 | 1 | 1 | 1 | 1 | 1 | 1 | 1 |
| D18S51 | 1 | 1 | 1 | 1 | 1 | 1 | 1 | 1 | 1 | 1 |
| D19S433 | 1 | 1 | 1 | 1 | 0 | 1 | 1 | 1 | 1 | 1 |
| D21S11 | 1 | 1 | 1 | 1 | 1 | 1 | 1 | 1 | 1 | 1 |
| D22S1045 | 0 | 1 | 1 | 1 | 0 | 0 | 1 | 0 | 1 | 1 |
| FGA | 1 | 1 | 1 | 1 | 1 | 1 | 1 | 1 | 1 | 1 |
| PENTA D | 0 | 0 | 0 | 0 | 1 | 1 | 0 | 1 | 1 | 1 |
| PENTA E | 0 | 0 | 0 | 0 | 1 | 1 | 0 | 1 | 1 | 1 |
| SE33 | 0 | 0 | 1 | 1 | 0 | 0 | 1 | 0 | 0 | 1 |
| TH01 | 1 | 1 | 1 | 1 | 1 | 1 | 1 | 1 | 1 | 1 |
| TPOX | 1 | 0 | 0 | 1 | 1 | 1 | 0 | 1 | 1 | 1 |
| vWA | 1 | 1 | 1 | 1 | 1 | 1 | 1 | 1 | 1 | 1 |
| SIBSHIP INDEX |  |  |  |  |  |  |  |  |  |  |
| Half Sibs $\mu$ | 5.1607 | 5.1954 | 5.5896 | 7.2936 | 5.1615 | 5.8591 | 5.5896 | 6.8599 | 7.5995 | 8.2835 |
| Half Sibs $\sigma$ | 1.4984 | 1.5798 | 1.6853 | 1.8474 | 1.5123 | 1.6044 | 1.6853 | 1.8115 | 1.862 | 1.9949 |
| Full Sibs $\mu$ | 6.7984 | 7.038 | 7.6848 | 9.7815 | 6.8367 | 7.7302 | 7.6848 | 9.3273 | 10.1427 | 11.2317 |
| Full Sibs $\sigma$ | 1.274 | 1.3473 | 1.4439 | 1.5818 | 1.2844 | 1.3655 | 1.4439 | 1.5467 | 1.5876 | 1.711 |
| Overlap | 0.55 | 0.53 | 0.5 | 0.47 | 0.55 | 0.53 | 0.5 | 0.46 | 0.46 | 0.42 |
| Sensitivity at cut off =1 | 0.99999995 | 0.99999991 | 0.99999995 | 1 | 0.99999995 | 0.999999993 | 0.99999995 | 1 | 1 | 1 |
| Specificity at cut off =1 | 0.00028644 | 0.00050351 | 0.00045565 | 0.00003940 | 0.0003212 | 0.00013017 | 0.00045565 | 0.00007628 | 0.00002240 | 0.00001646 |
| Crossing point | 5.8602 | 6.0088 | 6.5408 | 8.4536 | 5.8814 | 6.4841 | 6.5408 | 8.0134 | 8.789 | 9.6944 |
| Equal Error Rate point | 6.0458 | 6.1899 | 6.718 | 8.6339 | 6.0673 | 6.8699 | 6.718 | 8.1909 | 8.9723 | 9.8706 |
| Sensitivity=Specificity | 0.72263656 | 0.73549397 | 0.74842939 | 0.7659258 | 0.7254111 | 0.73566106 | 0.74842939 | 0.76875289 | 0.766952062 | 0.78685027 |
| SIBSHIP INDEX CORRECTED BY MATERNAL GENOTYPE |  |  |  |  |  |  |  |  |  |  |
| Half Sibs $\mu$ | -5.2462 | -5.4644 | -5.9402 | -7.5101 | -5.2855 | -5.9602 | -5.9402 | -7.2219 | -7.8282 | -8.6338 |
| Half Sibs $\sigma$ | 2.1202 | 2.1091 | 2.1633 | 2.4855 | 2.119 | 2.2549 | 2.1633 | 2.4496 | 2.5554 | 2.6697 |
| Full Sibs $\mu$ | 3.2970 | 3.6862 | 4.1663 | 4.9699 | 3.3539 | 3.7727 | 4.1663 | 4.9717 | 5.0939 | 5.9344 |
| Full Sibs $\sigma$ | 3.2202 | 3.3309 | 3.5084 | 3.9057 | 3.23339 | 3.4424 | 3.5084 | 3.8448 | 3.9641 | 4.2201 |
| Overlap | 0.1067 | 0.09 | 0.07 | 0.05 | 0.10 | 0.09 | 0.07 | 0.05 | 0.05 | 0.03 |
| Sensitivity at cut off =1 | 0.84705376 | 0.86578283 | 0.88249009 | 0.89839977 | 0.85015523 | 0.86344659 | 0.8824009 | 0.90201327 | 0.90060878 | 0.92017107 |
| Specificity at cut off =1 | 0.99332706 | 0.99521291 | 0.99698222 | 0.99874282 | 0.99369068 | 0.99589433 | 0.99698222 | 0.99840176 | 0.99890594 | 0.99938967 |
| Crossing point | -1.5259 | -1.572 | -1.7285 | -2.3097 | -1.5357 | -1.7756 | -1.7285 | -2.1328 | -2.4231 | -2.6387 |
| Equal Error Rate point | -1.8544 | -1.9167 | -2.0853 | -2.6567 | -1.8656 | -2.1081 | -2.0853 | -2.4765 | -2.7632 | -2.9888 |
| Sensitivity=Specificity | 0.94517378 | 0.953725 | 0.96261791 | 0.97457297 | 0.94673241 | 0.95621262 | 0.96261791 | 0.97364083 | 0.97626578 | 0.98276229 |

When SI algorithm is used and cutoff=1 is chosen as threshold, practically all Half-Sibs couples are declared as Full-Sibs; using cutoff=10^EERP^, 25% of couples are mis-classified.

When SICMG algorithm is used and cutoff=1 is chosen as threshold, less than 1% of Half-Sibs are declared Full-Sibs and about 12% of Full-Sibs are declared Half-Sibs; on the contrary, using cutoff=10^EERP^, about 3% of couples are misclassified; eight fold lower respect to SI algorithm. We defined “Core” the set of all 24 STRs showed in Table 4. Tables 5 and 6 show the performances obtained adding unusual STRs to the “Core” using SI and SICMG, respectively, and EERP as cutoff. Again, it is possible to observe the same phenomenon showed in Table 3: the Youden’s Index increases, reaches a maximum and then decreases.

**Table 5:**
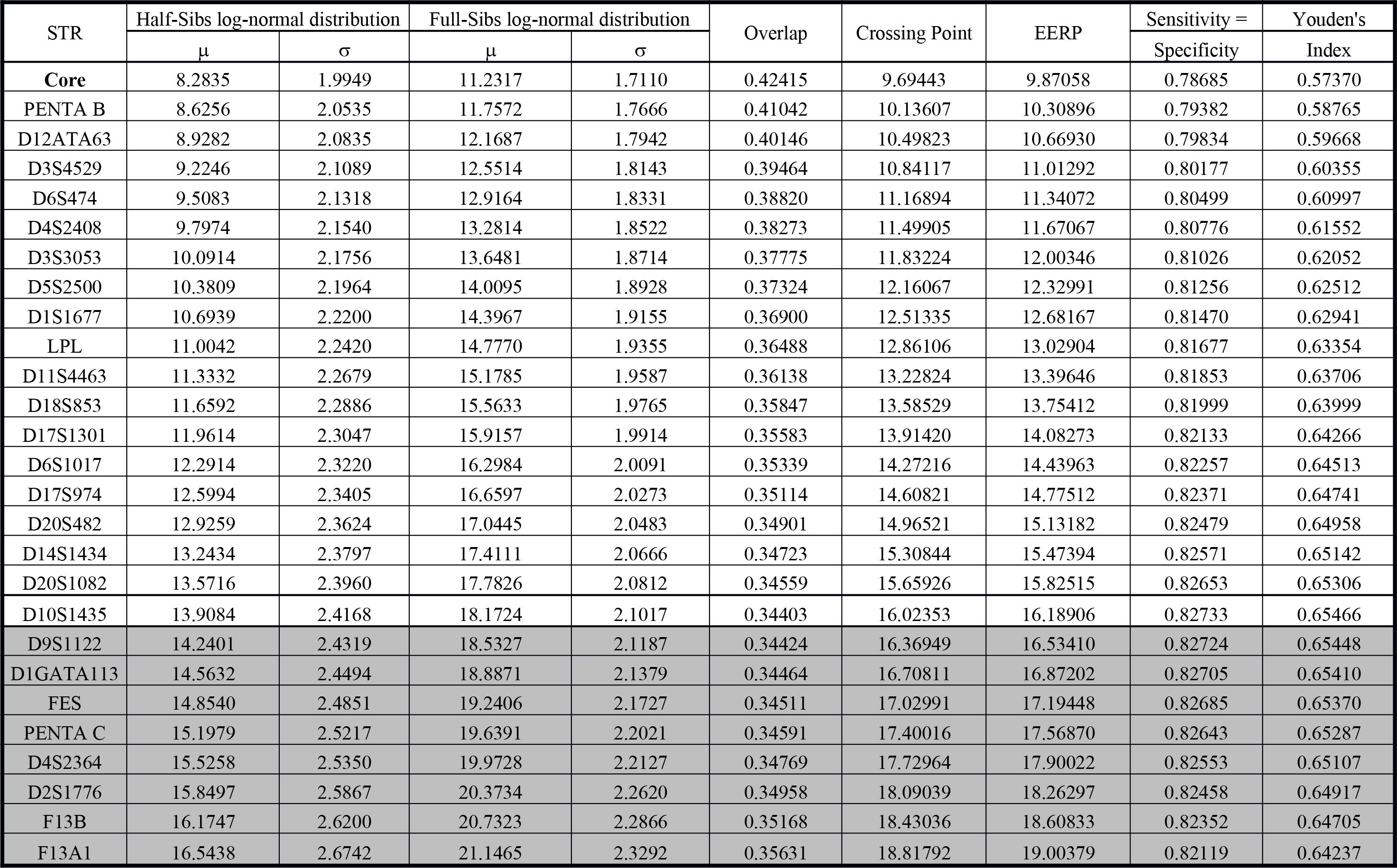
Performances of SI algorithm adding STRs to the Core of commercial used STR. In gray background STRs that worsen the performances.

**Table 6:**
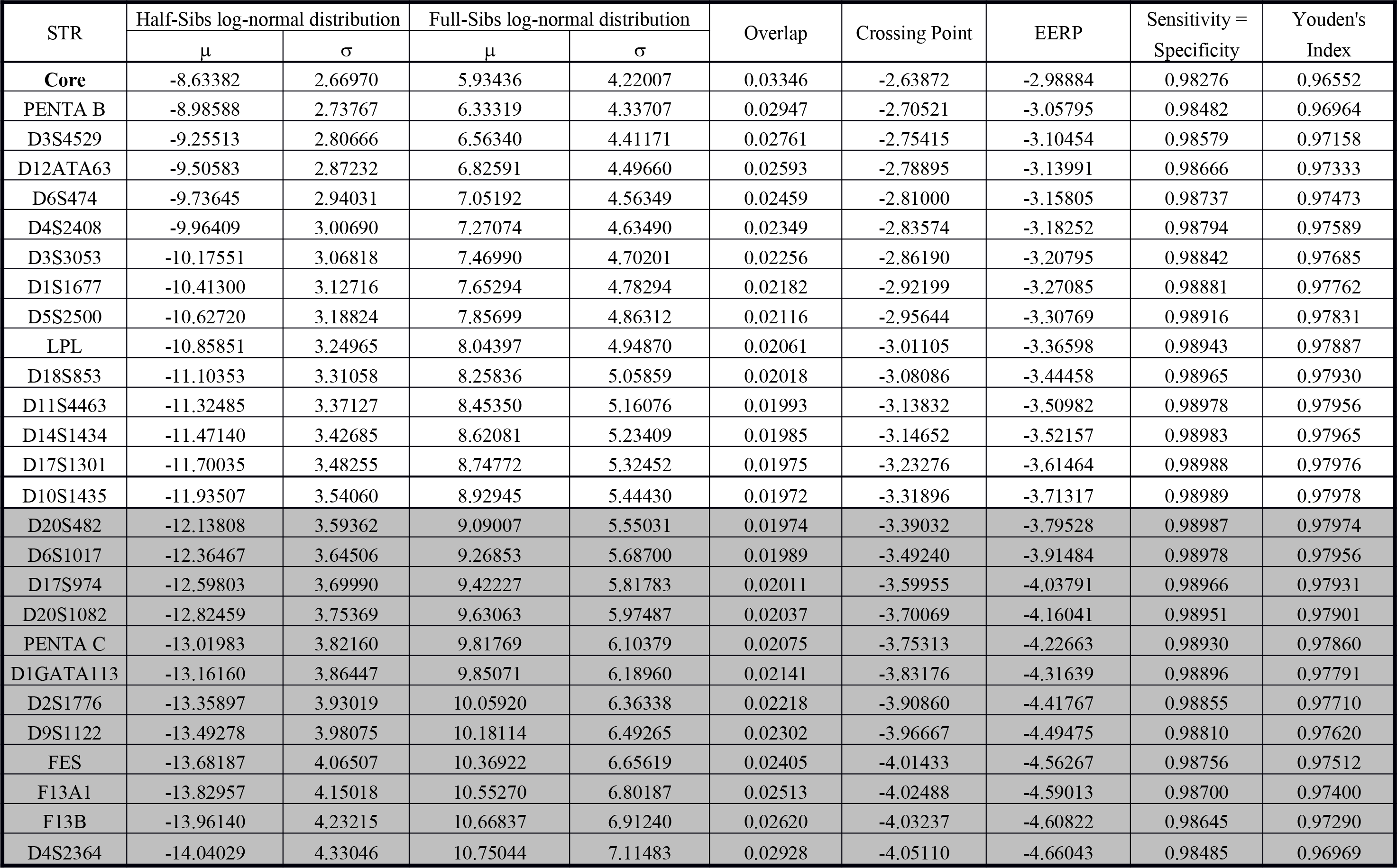
Performances of SICMG algorithm adding STRs to the Core of commercial used STR. In gray background STRs that worsen the performances.

| STR | Half-Sibs log-normal distribution |  | Full-Sibs log-normal distribution |  | Overlap | Crossing Point | EERP | Sensitivity =<br>Specificity | Youden's<br>Index |
| --- | --- | --- | --- | --- | --- | --- | --- | --- | --- |
| | $\mu$ | $\sigma$ | $\mu$ | $\sigma$ | | | | | |
| **Core** | -8.63382 | 2.66970 | 5.93436 | 4.22007 | 0.03346 | -2.63872 | -2.98884 | 0.98276 | 0.96552 |
| PENTA B | -8.98588 | 2.73767 | 6.33319 | 4.33707 | 0.02947 | -2.70521 | -3.05795 | 0.98482 | 0.96964 |
| D3S4529 | -9.25513 | 2.80666 | 6.56340 | 4.41171 | 0.02761 | -2.75415 | -3.10454 | 0.98579 | 0.97158 |
| D12ATA63 | -9.50583 | 2.87232 | 6.82591 | 4.49660 | 0.02593 | -2.78895 | -3.13991 | 0.98666 | 0.97333 |
| D6S474 | -9.73645 | 2.94031 | 7.05192 | 4.56349 | 0.02459 | -2.81000 | -3.15805 | 0.98737 | 0.97473 |
| D4S2408 | -9.96409 | 3.00690 | 7.27074 | 4.63490 | 0.02349 | -2.83574 | -3.18252 | 0.98794 | 0.97589 |
| D3S3053 | -10.17551 | 3.06818 | 7.46990 | 4.70201 | 0.02256 | -2.86190 | -3.20795 | 0.98842 | 0.97685 |
| D1S1677 | -10.41300 | 3.12716 | 7.65294 | 4.78294 | 0.02182 | -2.92199 | -3.27085 | 0.98881 | 0.97762 |
| D5S2500 | -10.62720 | 3.18824 | 7.85699 | 4.86312 | 0.02116 | -2.95644 | -3.30769 | 0.98916 | 0.97831 |
| LPL | -10.85851 | 3.24965 | 8.04397 | 4.94870 | 0.02061 | -3.01105 | -3.36598 | 0.98943 | 0.97887 |
| D18S853 | -11.10353 | 3.31058 | 8.25836 | 5.05859 | 0.02018 | -3.08086 | -3.44458 | 0.98965 | 0.97930 |
| D11S4463 | -11.32485 | 3.37127 | 8.45350 | 5.16076 | 0.01993 | -3.13832 | -3.50982 | 0.98978 | 0.97956 |
| D14S1434 | -11.47140 | 3.42685 | 8.62081 | 5.23409 | 0.01985 | -3.14652 | -3.52157 | 0.98983 | 0.97965 |
| D17S1301 | -11.70035 | 3.48255 | 8.74772 | 5.32452 | 0.01975 | -3.23276 | -3.61464 | 0.98988 | 0.97976 |
| D10S1435 | -11.93507 | 3.54060 | 8.92945 | 5.44430 | 0.01972 | -3.31896 | -3.71317 | 0.98989 | 0.97978 |
| D20S482 | -12.13808 | 3.59362 | 9.09007 | 5.55031 | 0.01974 | -3.39032 | -3.79528 | 0.98987 | 0.97974 |
| D6S1017 | -12.36467 | 3.64506 | 9.26853 | 5.68700 | 0.01989 | -3.49240 | -3.91484 | 0.98978 | 0.97956 |
| D17S974 | -12.59803 | 3.69990 | 9.42227 | 5.81783 | 0.02011 | -3.59955 | -4.03791 | 0.98966 | 0.97931 |
| D20S1082 | -12.82459 | 3.75369 | 9.63063 | 5.97487 | 0.02037 | -3.70069 | -4.16041 | 0.98951 | 0.97901 |
| PENTA C | -13.01983 | 3.82160 | 9.81769 | 6.10379 | 0.02075 | -3.75313 | -4.22663 | 0.98930 | 0.97860 |
| D1GATA113 | -13.16160 | 3.86447 | 9.85071 | 6.18960 | 0.02141 | -3.83176 | -4.31639 | 0.98896 | 0.97791 |
| D2S1776 | -13.35897 | 3.93019 | 10.05920 | 6.36338 | 0.02218 | -3.90860 | -4.41767 | 0.98855 | 0.97710 |
| D9S1122 | -13.49278 | 3.98075 | 10.18114 | 6.49265 | 0.02302 | -3.96667 | -4.49475 | 0.98810 | 0.97620 |
| FES | -13.68187 | 4.06507 | 10.36922 | 6.65619 | 0.02405 | -4.01433 | -4.56267 | 0.98756 | 0.97512 |
| F13A1 | -13.82957 | 4.15018 | 10.55270 | 6.80187 | 0.02513 | -4.02488 | -4.59013 | 0.98700 | 0.97400 |
| F13B | -13.96140 | 4.23215 | 10.66837 | 6.91240 | 0.02620 | -4.03237 | -4.60822 | 0.98645 | 0.97290 |
| D4S2364 | -14.04029 | 4.33046 | 10.75044 | 7.11483 | 0.02928 | -4.05110 | -4.66043 | 0.98485 | 0.96969 |

Table 5 shows that using SI algorithm and adding 18 STRs to the Core set the misclassified couples decrease from about 21% to 17%; the addition of other 8 STRs only succeeds in worsening the performances.

Similarly, Table 6 shows that using SICMG algorithm the maximum Youden’s Index is reached adding a set of 14 STRs but there is a gain of less than 1% in performances.

Anyway, there is a set of 8 STRs that are useless in both cases (D1GATA113, D2S1776, D4S2364, D9S1122, F13A1, F13B, FES and PENTA C).

In conclusion, it is impossible to discriminate Full-Sibs from Half-Sibs using usually SI index and cut-off; it is possible to greatly improve the performances taking into account the Maternal genotype and the Equivalent Error Rate Point, taking in mind that also in this condition 1-2% of couples will be misclassified.

